# Deep generalizable prediction of RNA secondary structure via base pair motif energy

**DOI:** 10.1101/2024.10.22.619430

**Authors:** Heqin Zhu, Fenghe Tang, Quan Quan, Ke Chen, Peng Xiong, S. Kevin Zhou

## Abstract

Deep learning methods have demonstrated great performance for RNA secondary structure prediction. However, generalizability is a common unsolved issue on unseen out-of-distribution RNA families, which hinders further improvement of the accuracy and robustness of deep learning methods. Here we construct a base pair motif library that enumerates the complete space of the locally adjacent three-neighbor base pair and records the thermodynamic energy of corresponding base pair motifs through *de novo* modeling of tertiary structures, and we further develop a deep learning approach for RNA secondary structure prediction, named BPfold, which learns relationship between RNA sequence and the energy map of base pair motif. Experiments on sequence-wise and family-wise datasets have demonstrated the great superiority of BPfold compared to other state-of-the-art approaches in accuracy and generalizability. We hope this work contributes to integrating physical priors and deep learning methods for the further discovery of RNA structures and functionalities.

## 1 Introduction

RNA secondary structure plays vital roles in modeling the RNA tertiary structure through base pairing interactions [1], demonstrating the remarkable versatility and functional mechanisms of RNA in biological systems and cellular processes, such as catalytic functionality [2, 3], regulatory functions [4], and intron splicing events [5]. Generally, RNA secondary structure forms a sequential of stem and loop regions, with stem regions composed of consecutive paired nucleotide bases [6, 7] and loop regions composed of unpaired bases. Furthermore, loop regions exhibit various structure motifs stabilized by non-canonical base pairs and other polar interactions, such as tetra loops, kissing-loops, kink turn, and G-quadruplex [8].

Discovering the secondary structure of RNA is important and necessary for modeling the tertiary structure and further exploring the potentialities of interactions between RNA structures and other biomolecules, such as proteins and ligands, which is crucial for drug design and RNA-based therapies [9, 10]. As the field of RNA research continues to expand, so does the need for precise and reliable detection of RNA secondary structures. Chemical probing techniques, such as SHAPE (Selective 2’-Hydroxyl Acylation analyzed by Primer Extension) [11], provide a way to infer the secondary structure of RNA molecules by selectively probing the reactivity of the RNA nucleotides. Advances in computational methods that predict RNA secondary structures from sequence data alone have greatly improved the effectiveness and efficiency for modeling RNA secondary structures, which enhances our fundamental knowledge of RNA biology and paves the way for innovative applications in medicine, biotechnology, and beyond [12].

During the past three decades, various computational methods have been developed to predict RNA secondary structures, such as comparative sequence analysis and thermodynamic models. Comparative sequence analysis [13–16] predicts structures by searching for homologous sequences, which is effective and accurate when the target sequence is hit in the homologous sequences database. However, the number of known RNA families is a few thousand in Rfam [17, 18], resulting in poor generalizability and accuracy of the comparative methods for unknown sequences. Thermodynamic models [19–30] aim at finding the best structure from thermodynamically stable candidates that are selected through free energy minimization. These methods assign each structure with a score, with parameters obtained from experiments, such as Vienna RNAfold [23, 31], RNAstructure [24, 32], and EternaFold [25]). Also, CONTRAfold, ContextFold, and EternaFold can be categorized into shallow machine learning (ML) methods. These non-ML methods and shallow learning methods are effective in predicting simple target secondary structures that only contain nested base pairs, while they have problems dealing with complex structures such as non-nested pairs (pseudoknots), and cannot predict non-canonical base pairs compared to end-to-end deep learning (DL) methods [33].

In recent years, deep learning methods [34–39] raise researchers’ significant attention, greatly boost the speed of prediction, and achieve high accuracy. As data-driven approaches, deep learning methods utilize the benefit of big data and learn the implicit features and intrinsic of data distribution in hidden space via deep neural networks. Once the neural network has learned to build the mapping and the relationship between input data (RNA sequence) and output data (RNA secondary structure), it can predict the secondary structure of an unknown arbitrary input RNA sequence. For instance, Singh et al. [36] propose SPOT-RNA, an ensemble of two-dimensional deep neural networks equipped with transfer learning on a high-quality dataset. Fu et al. [35] develop a U-shaped fully convolutional image-to-image network, named UFold, which converts an RNA sequence into an image-like representation to predict RNA secondary structure. Sato et al. [39] propose MXfold2 that learns RNA folding scores by integrating Turner’s nearest-neighbor free energy parameters into deep neural networks.

Although existing deep learning methods behave well on currently known test datasets, their performances degrade rapidly as sequence similarity decreases in situations of unseen RNA families and data distributions compared to non-ML methods [33, 40, 41], which indicates poor generalizability and the possibility of overfitting on training datasets. To mitigate this, researchers resort to integrating auxiliary information into deep learning models. For example, UFold [35] uses an alternative representation of the RNA sequence derived from CDPfold [42] to enhance the relationship between the input RNA sequence and thermodynamic prior of base pairs, SPOT-RNA [36] takes full advantage of evolutionary information, and MXfold2 [39] regularizes the learning of the model with a penalty loss on the folding score for deviating too far from thermodynamic estimation. With the help of auxiliary information, these methods have made some progress in improving prediction accuracy. However, a general data insufficiency and low data quality problem plaques RNA structure prediction, including secondary structure prediction. Unlike protein structure prediction, which possesses a sufficiently large number of high-quality data to represent the underlying distribution, which guarantees the effectiveness of DL methods such as AlphaFold [43, 44], the number, quality, and coverage of available RNA structure data are relatively very low [45]. Therefore, for RNA secondary structure prediction, how to develop a reliable DL model under such data insufficiency and further deal with out-of-distribution samples is an unsolved problem, hindering further improvements in the accuracy and generalizability of DL learning models.

It is known that enriching data at the secondary structure level is quite hard for both experimental and computational methods. Instead, it is more computationally efficient to predict the tertiary structure of short-sequence RNA motifs. Luckily, RNA secondary structure is mainly dependent on the structure motifs [46]. Motivated by these and different from previous attempts that integrate knowledge prior into data-driven models incompletely, we propose to leverage the local short-distance interactions of base pairs and enumerate the whole space of adjacent neighboring patterns of all canonical base pairs, named as base pair motif, aiming at enriching data at the base-pair level completely.

In this paper, we propose BPfold, a deep learning model integrated with thermodynamic energy from the complete space of the upstream and downstream of three-neighbor base pair motifs for predicting RNA secondary structures. BPfold comprises two key components:

1. *Base pair motif energy*. A base pair motif is a canonical base pair (i.e., A-U, U-A, G-C, C-G, G-U, and U-G) together with its local spatially adjacent bases, which dominates the local structure of the base pair. We explore the entire space of the base pair motifs of *r* neighbors and compute their energy by *de novo* modeling the tertiary structure of the base pair motif. After storing the computed tertiary structures and corresponding energy items in the motif library, we can quickly obtain the base pair motif energy for any base pair of any arbitrary input RNA sequence. Thereby, this auxiliary input energy fully covers the data distribution at the base-pair level, mitigating the current insufficient database, and eliminating the major hurdle posed by the generalizability of *de novo* DL models.
2. *Base pair attention*. In the BPfold neural network, we elaborately design a base pair attention block, which combines transformer [47] and convolution [48] layers and enables information integration between RNA sequence and base pair motif energy. The base pair attention block aggregates the attention map of the RNA sequence and the base pair motif energy to effectively learn the base pair knowledge from the RNA sequence. BPfold takes advantage of the deep learning approach, predicts the accurate RNA secondary structure in seconds, and has great generalizability that accounts for the learned knowledge of thermodynamic energy.

We conduct sequence-wise and family-wise cross-validation experiments on multiple benchmark datasets to evaluate the accuracy and generalizability of BPfold. ArchiveII [49] (3,966 RNAs) and bpRNA-TS0 [50] (1,305 RNAs) are sequence-wise datasets while Rfam12.3-14.10 [17, 18] (10,791 RNAs) contains cross-family RNA sequences and PDB [51] (116 RNAs) consists of high-quality experimentally validated RNA structures. Quantitative and qualitative results demonstrate the superiority of the proposed BPfold in accuracy and generalizability against other learning-based methods and non-learning methods. We expect this work will take a meaningful step toward fast and robust prediction of RNA secondary structures.

## 2 Results

### 2.1 Overview of the BPfold approach

In this work, we present BPfold (Fig. 1a), a deep learning approach integrated with thermodynamic energy for RNA secondary structure prediction. Aiming at improving the generalizability and accuracy of the deep learning-based model, we compute the base pair motif energy and design a base pair attention neural network block. As demonstrated in Fig. 2, a base pair motif is a canonical base pair (i.e., A-U, U-A, G-C, C-G, G-U, and U-G) together with their neighboring bases. We compute the *de novo* RNA tertiary structure and obtain its thermodynamic energy of the complete space of *r*-neighbor base pair motif to mitigate the insufficient coverage of base pair in existing datasets. Furthermore, as illustrated in Fig. 1b, we equip BPfold with a custom-designed base pair attention block (described in Section 4.2), which applies an attention mechanism to the base pair motif energy (described in Section 4.1) and RNA sequence feature to perfectly learn representative knowledge from RNA sequence and thermodynamic energy.

**Fig. 1:**
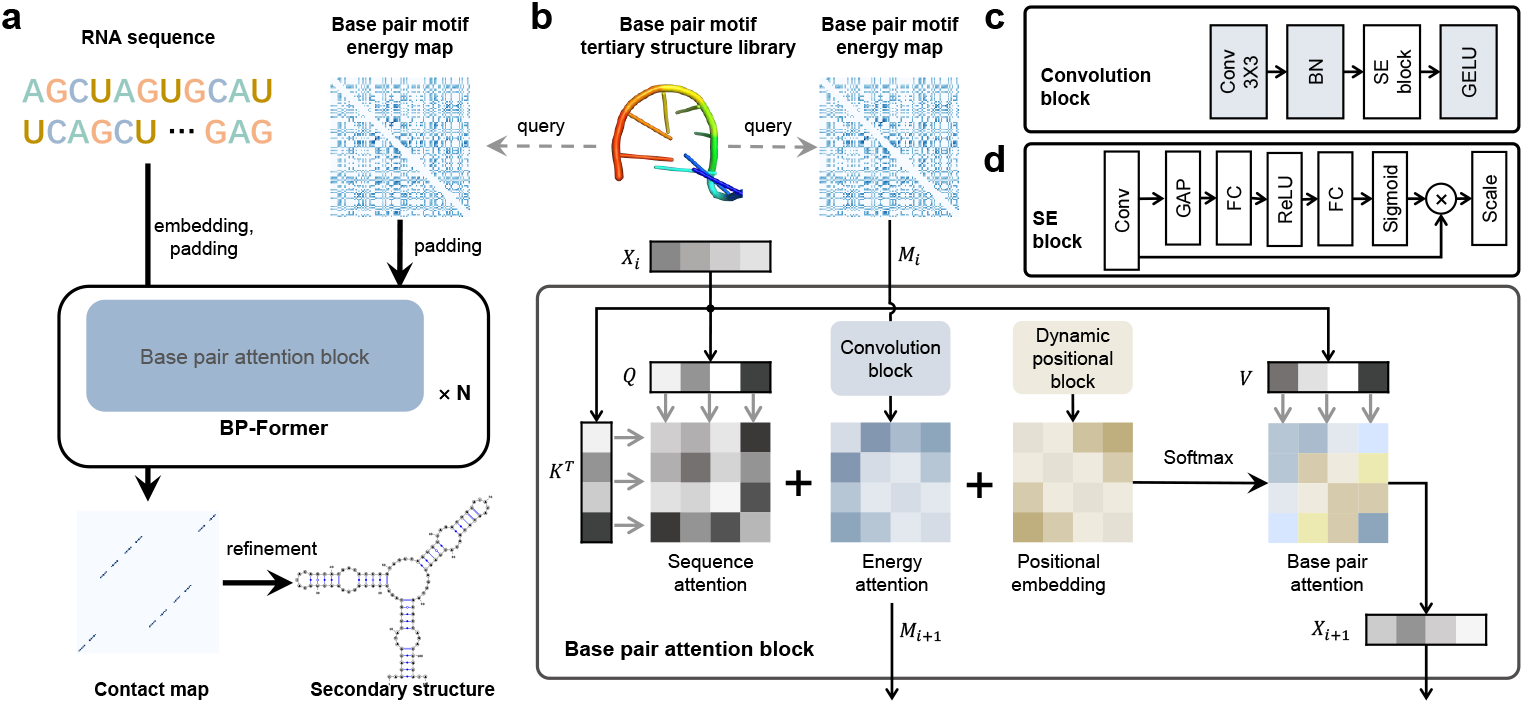
Overview of our BPfold approach for modeling RNA secondary structures. **a** BPfold takes RNA sequence and corresponding base pair motif energy map generated from base pair motif library as inputs, consisting of transformer blocks with designed base pair attention, and outputs contact map. After applying physical constraints to the contact map in refinement procedures, we obtain the final predicted secondary structure. **b** The detailed structure of the proposed base pair attention block which jointly fuses the sequence features *X*_*i*_, *X*_*i*+1_ and energy matrix features *M*_*i*_, *M*_*i*+1_ for enhanced learning of base pair interactions. *Q, K, V* represent query, key, and value matrix when computing self-attention. **c** The detailed structure of convolution block in base pair attention block. **d** The detailed structure of squeeze & excitation (SE) block in convolution block.

**Fig. 2:**
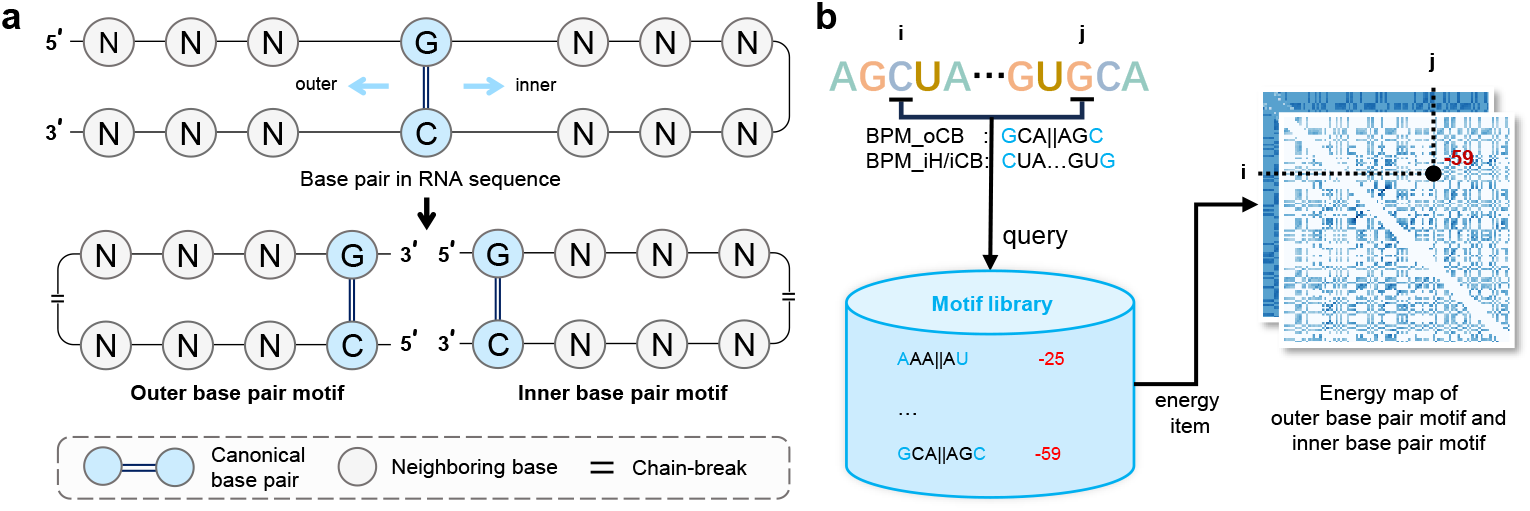
Diagram of generating outer/inner base pair motif from a canonical base pair and constructing energy maps of base pair motifs for an RNA sequence. **a** For any canonical base pair (i.e., A-U, U-A, G-C, C-G, G-U, and U-G) in an RNA sequence, the upstream and downstream three neighboring bases (denoted as N, an arbitrary base of A, U, G, or C) of the base pair form two base pair motifs, an inner base pair motif with neighboring bases extending to the middle of the RNA sequence, and an outer base pair motif with neighboring bases extending to both ends of the RNA sequence. Note that the inner base pair motif can be categorized into inner hairpin base pair motif and inner chain-break base pair motif (demonstrated in this figure) in accordance with the distance of the paired bases. **b** For any canonical base pair (*i, j*) from an RNA sequence of *L* nucleotides, we firstly find the corresponding outer/inner base pair motifs of this base pair and then query the energy items in the base pair motif library, which forms the (*i, j*) element of the outer/inner energy maps in a shape of *L* × *L*.

### 2.2 Establishing the base pair motif library

As Fig. 2 shows, we define three categories of base pair motifs, namely (inner) hairpin base pair motif, inner chainbreak base pair motif and outer chainbreak base pair motif, which are denoted as BPM_iH_, BPM_iCB_ and BPM_oCB_, respectively. We build the base pair motif library by modeling the tertiary structures of all three-neighbor base pair motifs and storing corresponding energy items in the motif library. Specifically, each tertiary structure of base pair motif is computed by our previous *de novo* RNA structure modeling method BRIQ [52], which employs Monte Carlo (MC) [53] algorithm to sample candidate RNA tertiary structures and evaluates the BRIQ energy score of each sampled tertiary structure. BRIQ energy score is a combined energy score of physical energy using density functional theory (DFT) and statistical energy calibrated by quantum mechanism, supplying a trade-off measurement of thermodynamic energy between computational speed and accuracy. Furthermore, each energy score of the base pair motif is normalized according to its sequence length and motif category. After obtaining all energy items, given an RNA sequence of length *L*, for any base *i* and base *j*, we built two energy maps in the shape of *L* × *L*. One for the outer base pair motif and the other for the inner base pair motif denoted as *M*^*µ*^ and *M*^*ν*^, respectively, which are used as input thermodynamic information by the BPfold neural network.

After constructing the base pair motif library, we make a detailed analysis of this library, which is displayed in Fig. 3. Firstly, we demonstrate the coverage of the base pair motif in current datasets. As Fig. 3a shows, the RNAStrAlign dataset contains adequate chainbreak base pair motifs with only 249 chainbreak motifs missing. However, as for the hairpin motifs, RNAStrAlign misses 24,576 motifs, weighting 32.3% of total 75,990 motifs, covering a small data distribution, which hinders the pattern recognition of hairpin base pair motifs for deep learning models (See the data coverage of the other four datasets and the intersection of base pair motifs from these datasets in Supplementary Fig. 1 and Supplementary Fig. 2, respectively). Furthermore, we store the intermediate results of the base pair motif energy map when BPfold predicts secondary structures from the third convolutional layer, and then apply t-SNE [54] decompression to these feature maps to visualize the latent embeddings of base pair energy maps from the six largest RNA families (accounting for approximate 90%) of ArchiveII dataset. As Fig. 3b demonstrates, BPfold learns discriminative embeddings of family-wise RNAs effectively, projecting the features of energy maps to a wide-range scattered latent vectors. In addition, we visualize the heatmap of the stored intermediate feature map of the base pair energy map from the third convolutional layer in

**Fig. 3:**
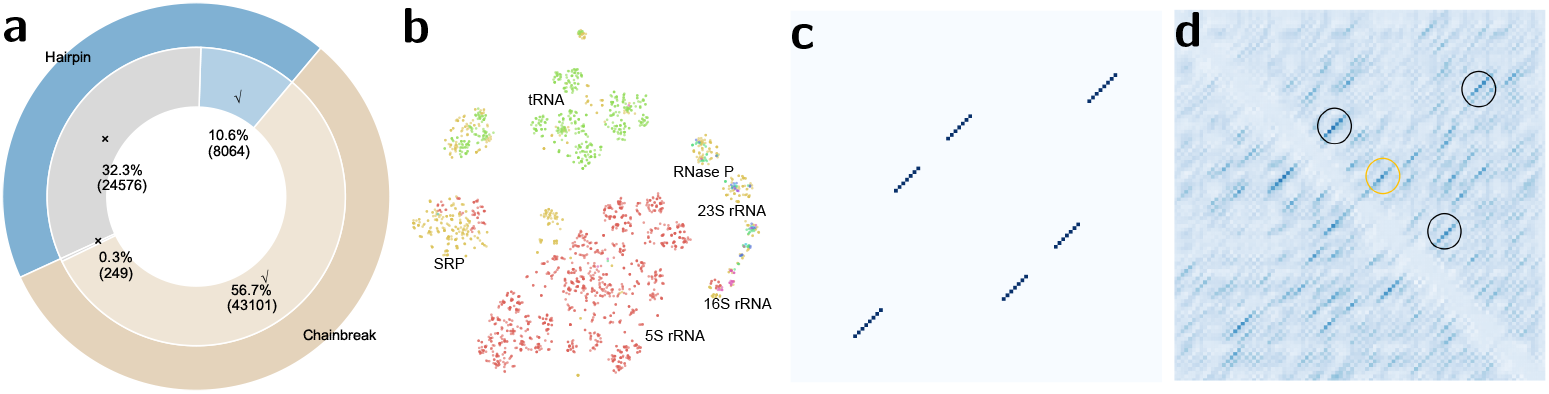
Analysis of base pair motif library. **a** Pie visualization of the data coverage of hairpin and chainbreak (including inner and outer) base pair motifs in current datasets, which are denoted as BPM_iH_, BPM_iCB_ and BPM_oCB_, respectively. The outer ring of each pie chart represents the hairpin and chainbreak distribution in the whole base pair motif library (75,990 motifs, blue for hairpin motifs, brown for chainbreak motifs). The inner ring represents the hairpin and chainbreak distribution in each dataset (light blue for hairpin motifs, light brown for chainbreak motifs, and grey for missing motifs). **b** t-SNE visualization of the latent feature map of base pair motif energy map at the third convolutional layer from various RNA families in ArchiveII dataset (n=3,966 RNAs). **c** Ground truth heatmap visualization of the secondary structure of an example RNA sequence. **d** Heatmap visualization of the extracted latent feature map of the same RNA sequence from subfig **c**. The corrected responses of base pair interactions are annotated in black circles.

Fig. 3d. Compared with the ground truth heatmap displayed in Fig. 3c, the feature map correctly captures the interactions of base pairs, annotated in black circles, which indicates that base pair motif energy assists the neural network with the interpretation of base pair interactions. Although there are false positive interactions annotated in yellow circles, these weak responses will be eliminated by subsequent transformer layers and refinement procedures. In view of the above analysis of the base pair motif library, we expect that our proposed base pair motif library that takes into account tertiary structures and energy items can positively contribute to any computational method.

### 2.3 Assessing the effectiveness of base pair motif energy

To investigate the effectiveness of the proposed base pair motif energy, which is the key contribution of BPfold, we conduct ablation studies to demonstrate the performances of BPfold (1) with and without base pair motif energy; (2) with one category of base pair motif energy. In this experiment, we train BPfold on training datasets of RNAStrAlign [55] and bpRNA-1m [50] under five different configurations of base pair motif energy: (1) with BPM (all); (2) without BPM; (3) with BPM_iH_; (4) with BPM_iCB_; (5) with BPM_oCB_, respectively, and evaluate them on family-wise dataset Rfam12.3-14.10 [17, 18].

As shown in Fig. 4a and Supplementary Tables 2, BPfold with all base pair motif energy achieves INF, F1, precision and recall of 0.694, 0.689, 0.660 and 0.741, respectively, behaving better than any other configurations under all metrics (∀ p value *<* 0.001 using one-sided t-test on F1 score, such as BPM_iCB_ with p value = 6.066^−17^), indicating that each category of base pair motif energy is essential for addressing the gaps in data distribution at the base-pair level regarding thermodynamic energy, significantly enhancing performance on unseen data and providing BPfold with improved generalizability and robustness. Fig. 4b and Supplementary Table 2 demonstrate the detailed results of BPfold with and without base pair motif energy on the Rfam12.3-14.10 dataset and its five specific RNA families with the most RNA sequences: Cobalamin (riboswitches that regulate adjacent genes), skipping-rope (RNA motifs likely function in translate as small RNAs), Twister-P1 (ribozymes), Cyclic di-GMP-II (riboswtiches that are common in species within the class Clostridia and the genus Deinococcus) and RAGATH-18 (self-cleaving ribozymes in bacteria). BPfold with base pair motif energy achieves much better performances than BPfold without base pair motif energy on these unseen RNA families, indicating that the thermodynamic energy greatly improves the generalizability and accuracy of deep learning models.

**Fig. 4:**
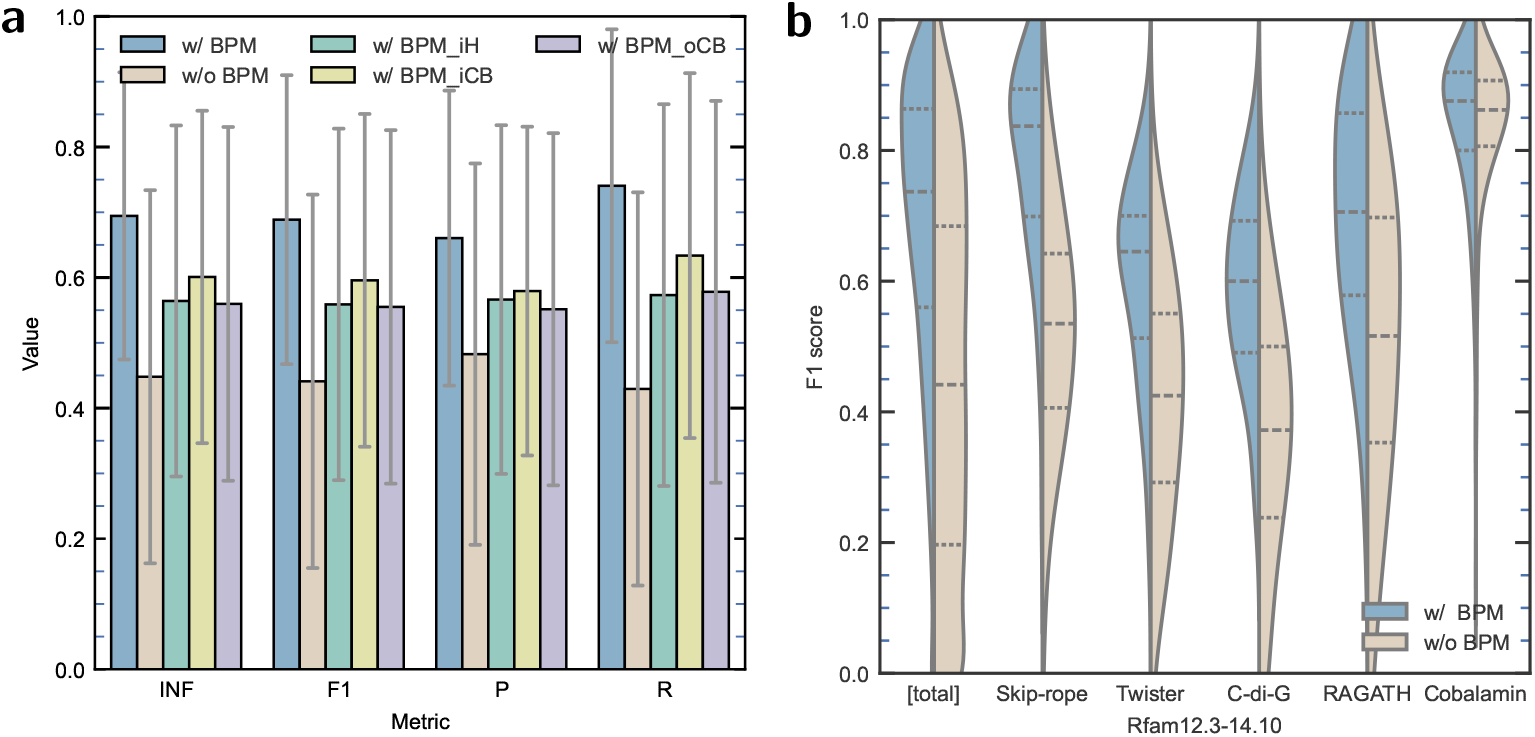
BPfold performance under different configurations. **a** Ablation study of BPfold under five configurations on family-wise dataset Rfam12.3-14.10 (n=10,791 RNAs): (1) with BPM (all); (2) without BPM; (3) with BPM_iH_; (4) with BPM_iCB_; (5) with BPM_oCB_. Data are presented as mean values ± SD. **b** Ablation study of BPfold with and without base pair motif energy on Rfam12.3-14.10 (n=10,791 RNAs) and its five specific RNA: Skip-rope (n=240 RNAs), Twister (n=236 RNAs), C-di-G (n=197 RNAs), RAGATH (n=160 RNAs), and Cobalamin (n=286 RNAs). The 25th percentiles, median, 75th percentiles are shown as dashed lines from bottom to top, while whiskers spanning the full data range of [0, 1]. With the integration of all base pair motifs, BPfold improves the prediction accuracy by a large gap on unseen RNA families.

Additionally, we make head-to-head comparisons of BPfold with BPM energy compared to the other four configurations on Rfam12.3-14.10. As shown in Supplementary Fig. 3, head-to-head analysis demonstrates that the accuracy of BPfold with (all) base pair motif energy is above that of any other configuration in the majority of sample points, indicating the advantage of base pair motif energy is much clearer on improving accuracy and generalizability of learning-based data-driven neural networks.

### 2.4 Evaluating BPfold on sequence-wise datasets

In this and the following subsection, we compare proposed BPfold with other nine state-of-the-art methods, including (1) Deep learning methods: SPOT-RNA (committed on Jun. 23, 2022) [36] and MXfold2 version 0.1.2 [39]; (2) Shallow learning methods: ContextFold version 1.0 [20], CONTRAfold version 2.02 [19] and EternaFold version 1.3.1 [25]; (3) Non-learning methods: Linearfold (committed on Aug. 29, 2022) [22], RNAfold in ViennaRNA package version 2.6.4 [23, 31], SimFold [26] in MultiRNAFold [56] package version 2.0 in RNAsoft package [57] and RNAstructure version 6.4 [24, 32]. The training datasets are the same for trainable models (i.e. BPfold, SPOT-RNA, MXfold2, CONTRAfold), namely RNAStrAlign [55] and bpRNA [50], except SPOT-RNA that employs PDB dataset [51] for transfer learning and can not be retrained because it does not disclose training module, while the other methods use default parameters. For a fair comparison with SPOT-RNA, we train BPfold in a manner of 5-fold cross-validation and apply early stopping to prevent over-fitting. All methods are evaluated on sequence-wise test datasets that contain distinguished sequences from training datasets and family-wise test datasets that consist of unseen RNA families as out-of-distribution validation under measurements of interaction network fidelity (INF) [58], F1-score, precision, and recall (sensitivity).

To evaluate the performance of our BPfold model on sequence-wise datasets, we report the results of BPfold on ArchiveII [49] test set and bpRNA-TS0 [50] test set, compared with the above deep learning methods, shallow learning method, and non-learning methods. Same as previous deep learning methods [35, 37, 39], for fair comparison and resource saving, BPfold is trained on RNA sequences with lengths no more than 600 nucleotides from RNAStrAlign [55] and bpRNA [50] datasets.

As Fig. 5a and Table 1 demonstrate, on bpRNA-TS0 dataset, non-ML methods achieve an average F1 score in the range of [0.507, 0.530] and shallow learning (SL) methods achieve an average F1 score in the range of [0.516, 0.547], dropping behind deep learning methods such as 0.575 of Mxfold2 and 0.625 of SPOT-RNA. With the base pair motif energy, BPfold further improves the performance and obtains an average F1 score of 0.658, leading ahead of other methods by a remarkable gap (∀ p value *<* 0.001 using one-sided t-test on F1 score, such as SPOT-RNA with p value = 8.988 × 10^−4^, MXfold2 with p value = 4.297 × 10−14, and ContextFold with p value = 3.718 × 10^−36^). Compared with the previous state-of-the-art method SPOT-RNA, BPfold achieves an about 5% increase in F1 score and INF metric. As for the ArchiveII dataset, Fig. 5b and Table 1 demonstrate similar rankings except ContextFold obtains the second place. BPfold also reaches the highest prediction accuracy, achieving an average F1 score of 0.820 and an INF of 0.823, significantly outperforming any other methods (∀ p value *<* 0.001 using one-sided t-test on F1 score, such as SPOT-RNA with p value = 5.662 × 10^−76^ and MXfold2 with p value = 1.119 × 10^−121^) except ContextFold (p value = 0.345 using one-sided t-test on F1 score). ContextFold obtains an F1 score of 0.818 and an INF of 0.820 on the ArchiveII dataset, slightly lower than that of BPfold. In general, BPfold predicts more accurate RNA secondary structures compared with other methods on these sequence-wise datasets from the same data distribution.

**Table 1:** Sequence-wise evaluation of three deep learning methods (BPfold, SPOT-RNA, and MXfold2), three shallow learning methods (ContextFold, CONTRAfold, and EternaFold), and non-ML methods (LinearFold, RNAfold, SimFold, and RNAs-tructure) on bpRNA-TS0 (n=1,305 RNAs) and ArchiveII (n=3,966 RNAs) datasets.

| Method | bpRNA-TS0 |  |  |  | ArchiveII |  |  |  |
| --- | --- | --- | --- | --- | --- | --- | --- | --- |
|  | INF | F1 | Precision | Recall | INF | F1 | Precision | Recall |
| BPfold | 0.670 | 0.658 | 0.599 | 0.770 | 0.823 | 0.820 | 0.818 | 0.834 |
| SPOT-RNA | 0.634 | 0.625 | 0.598 | 0.694 | 0.736 | 0.730 | 0.763 | 0.723 |
| MXfold2 | 0.587 | 0.575 | 0.521 | 0.682 | 0.711 | 0.709 | 0.697 | 0.728 |
| ContextFold | 0.526 | 0.516 | 0.477 | 0.599 | 0.820 | 0.818 | 0.824 | 0.820 |
| CONTRAfold | 0.557 | 0.547 | 0.515 | 0.625 | 0.597 | 0.594 | 0.612 | 0.588 |
| EternaFold | 0.553 | 0.539 | 0.475 | 0.663 | 0.601 | 0.599 | 0.573 | 0.636 |
| LinearFold | 0.539 | 0.530 | 0.539 | 0.582 | 0.610 | 0.606 | 0.629 | 0.605 |
| RNAfold | 0.522 | 0.508 | 0.446 | 0.631 | 0.579 | 0.577 | 0.551 | 0.613 |
| SimFold | 0.520 | 0.507 | 0.448 | 0.626 | 0.570 | 0.568 | 0.550 | 0.594 |
| RNAstructure | 0.520 | 0.507 | 0.446 | 0.628 | 0.575 | 0.573 | 0.548 | 0.607 |

**Fig. 5:**
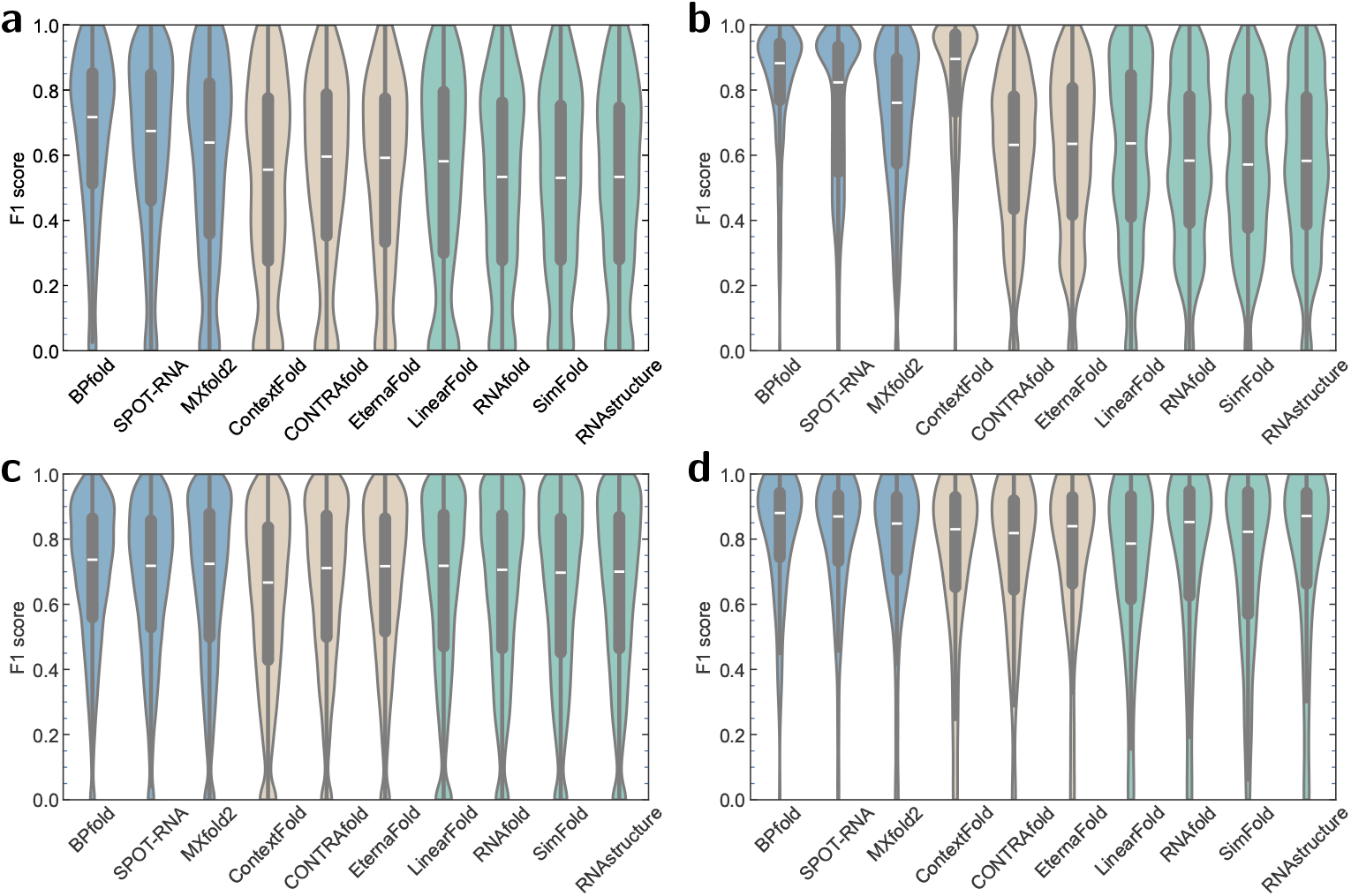
Performance comparison under macro-average F1 measurement of BPfold and nine other RNA secondary structure prediction methods. Deep learning methods, shallow learning methods, and non-learning methods are marked as blue, brown, and green, respectively. The median is marked as white, while the 25th and 75th percentiles are indicated by the bottom and top of black band, respectively. Each whisker spans the full data range of [0, 1]. **a** Sequence-wise dataset bpRNA-TS0 (n=1,305 RNAs). **b** Sequence-wise dataset ArchiveII (n=3,966 RNAs). **c** Family-wise dataset Rfam12.3-14.10 (n=10,791 RNAs). **d** High-quality dataset PDB (n=116 RNAs).

### 2.5 Evaluating BPfold on family-wise datasets

With the same configurations and training procedures, we further evaluate our BPfold model on family-wise datasets of unseen RNA families from out-of-distribution data to verify its model generalizability. Since the training set bpRNA contains RNA sequences from Rfam version 12.2, we collect Rfam12.3-14.10 dataset from Rfam [17, 18] database by retaining the newly added unseen families of version 14.10 from version 12.3, which contains 10,791 RNA sequences from 1,992 RNA families after removing similar sequences by CD-HIT-EST [59] with a threshold of 80%.

As Fig. 5c and Table 2 demonstrate, deep learning methods obtain satisfying performances even though the family-wise test dataset is out of distribution, such as Mxfold2 obtains F1 score of 0.664 on Rfam12.3-14.10 and SPOT-RNA achieves an F1 score of 0.672. As mentioned above, Mxfold2 takes advantage of integrating free energy minimization into the deep learning model and SPOT-RNA utilizes evolutionary information. However, these kinds of auxiliary information are not complete and bias-free, which hinders the overall performance of unseen data. In contrast, BPfold outperforms any other deep learning methods and non-DL methods (∀ p value *<* 0.001 using one-sided t-test on F1 score, such as SPOT-RNA with p value = 3.403 × 10^−8^) with the help of thermodynamic energy, exploring the complete space of 3-neighbor base pair motif and mitigating the out-of-distribution data at base-pair level, achieving the best F1 score of 0.689 and INF of 0.694. However, non-DL methods also achieve comparable performances with the help of thermodynamic parameters and physical laws, such as EternaFold and RNAstructure. We also evaluate the performances of the above methods on bpRNA-new, a subset of Rfam12.3-14.10, which contains 5,401 RNAs with a maximum sequence length of 439 nucleotides from Rfam version 12.3 to Rfam version 14.2. As Supplementary Table 3 shows, BPfold also wins first place on the bpRNA-new dataset, obtaining an F1 score of 0.647 and an INF of 0.655, indicating the great power of the proposed base pair motif for improving the generalizability in family-wise evaluation. It is worth mentioning that the non-ML method LinearFold achieves the second-best precision of 0.677 compared to the best precision of 0.678 and shallow learning method EternaFold which employs thermodynamic parameters wins the first place in recall of 0.746, which indicates the effectiveness of non-ML approaches in modeling RNAs from new families to some extent.

**Table 2:**
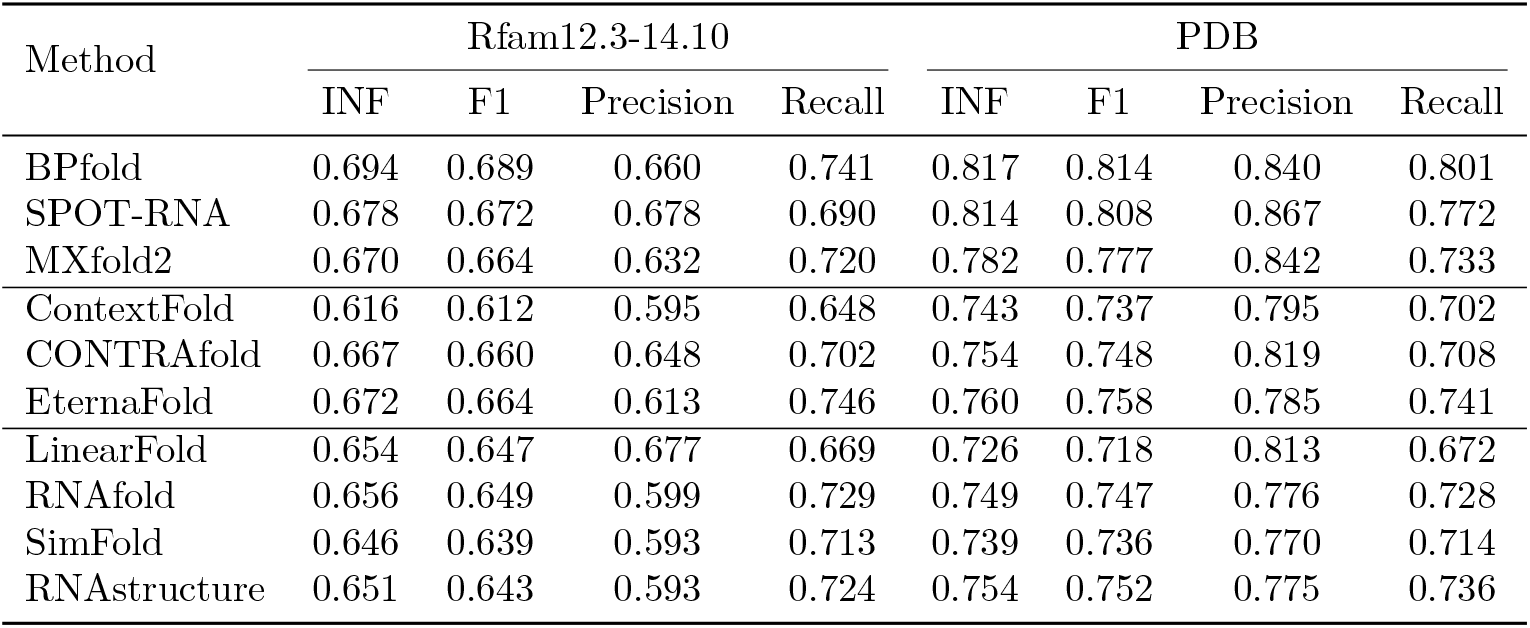
Family-wise evaluation of three deep learning methods (BPfold, SPOT-RNA, and MXfold2), three shallow learning methods (ContextFold, CONTRAfold, and EternaFold) and non-learning methods (LinearFold, RNAfold, SimFold, and RNAs-tructure) on Rfam12.3-14.10 (n=10,791 RNAs) and PDB (n=116 RNAs) datasets.

| Method | Rfam12.3-14.10 |  |  |  | PDB |  |  |  |
| --- | --- | --- | --- | --- | --- | --- | --- | --- |
|  | INF | F1 | Precision | Recall | INF | F1 | Precision | Recall |
| BPfold | 0.694 | 0.689 | 0.660 | 0.741 | 0.817 | 0.814 | 0.840 | 0.801 |
| SPOT-RNA | 0.678 | 0.672 | 0.678 | 0.690 | 0.814 | 0.808 | 0.867 | 0.772 |
| MXfold2 | 0.670 | 0.664 | 0.632 | 0.720 | 0.782 | 0.777 | 0.842 | 0.733 |
| ContextFold | 0.616 | 0.612 | 0.595 | 0.648 | 0.743 | 0.737 | 0.795 | 0.702 |
| CONTRAfold | 0.667 | 0.660 | 0.648 | 0.702 | 0.754 | 0.748 | 0.819 | 0.708 |
| EternaFold | 0.672 | 0.664 | 0.613 | 0.746 | 0.760 | 0.758 | 0.785 | 0.741 |
| LinearFold | 0.654 | 0.647 | 0.677 | 0.669 | 0.726 | 0.718 | 0.813 | 0.672 |
| RNAfold | 0.656 | 0.649 | 0.599 | 0.729 | 0.749 | 0.747 | 0.776 | 0.728 |
| SimFold | 0.646 | 0.639 | 0.593 | 0.713 | 0.739 | 0.736 | 0.770 | 0.714 |
| RNAstructure | 0.651 | 0.643 | 0.593 | 0.724 | 0.754 | 0.752 | 0.775 | 0.736 |

Furthermore, we evaluate BPfold on PDB, a widely used benchmark dataset that contains high-resolution RNA X-ray tertiary structures. We divide this set into three test sets as SPOT-RNA does, namely TS1, TS2, and TS3, which contain 60, 38, and 18 sequences, respectively. As Fig. 5d, Table 2 and Supplementary Table 3, 4 demonstrate the F1 score, precision, and recall metrics of canonical pairs (i.e., A-U, U-A, G-C, C-G, G-U and U-G), BPfold achieves an F1 score of 0.814 and an INF of 0.817, behaves better than any other methods, demonstrating high prediction accuracy and strong generalizability in detecting dense canonical pairs of RNA secondary structures on this high-quality experimentally validated dataset. While the F1 score of BPfold is not significantly better than SPOT-RNA (p value = 0.407 using one-sided t-test on F1 score) and MXfold2 (p value = 0.086), BPfold predicts more accurate structures than other methods, such as ContextFold (p value = 0.005), CONTRAfold (p value = 0.010), EternaFold (p value = 0.026). SPOT-RNA utilizes the PDB dataset for transfer learning, which could explain why the difference in F1 score between BPfold and SPOT-RNA on the PDB dataset is minimal.

### 2.6 Visualizing the performance of BPfold

To demonstrate the efficiency of RNA secondary structures prediction methods and evaluate the prediction time at the inference stage, we conduct experiments on a selected dataset that contains 174 RNAs with lengths varying from 60 to 1,851 nucleotides uniformly. To decrease the influence of randomness, we repeat 10 times for each RNA sequence. We select deep learning methods SPOT-RNA and MXfold2, shallowing learning method CONTRAfold, and non-learning method LinearFold for comparison because it is particularly hard and slow for some methods to predict long RNA sequences such as RNAstructure. The results demonstrated in Supplementary

Fig. 4 show that BPfold predicts RNA secondary structures within 10 seconds for RNAs no longer than 1,000 nucleotides, and within 40 seconds for RNAs no longer than 1,851 nucleotides, which is comparable with MXfold2 and CONTRAfold. SPOT-RNA takes several times longer time for predicting long sequences while LinearFold is the fastest method that predicts secondary structures within 1 second for RNAs no longer than 1,851.

Apart from the above quantitative experimental analysis, we also provide qualitative visualization of the RNA secondary structures predicted by BPfold to verify the detailed interactions of each nucleotide. To achieve this, we utilize the RNA visualization tool VARNA [62] to draw the figure, which takes various formats of RNA secondary structures, such as dot-bracket notation, bpseq, and ct format. For comparison, we also show the structures of native annotations (ground truth structures) and other two excellent methods, the deep-learning method SPOT-RNA and the traditional method CONTRAfold. As Fig. 6 visualizes, each column displays the structures of one method and each row displays one example. In these samples, structures predicted by BPfold are similar to native structures, and more accurate and robust than other methods. Specifically, Fig. 6a shows the superiority of BPfold in precision on an example of brucella abortus S19 signal recognition particle RNA, with 97% precision and 94% recall, respectively. Meanwhile, Fig. 6b displays an example of the deltaJ-delta-K domain of EMCV IRES [60] from PDB [51] dataset (PDB ID=2NC1) in radiate style, where lines with blue solid circle denote non-canonical pairs. As it shows, BPfold has the ability to predict non-canonical pairs, outperforming other methods, with 100% precision and 93% recall, respectively. Fig. 6c displays the structures of the lariat capping ribozyme [61] from PDB [51] dataset (PDB ID=4P95) in a circular style. This RNA sequence is a long sequence with 192 nucleotides and dense interactions. BPfold successfully models the long-range connections and predicts the most accurate structure than other methods, with 93% precision and 96% recall, respectively. Furthermore, we display the effect of applying removing isolated base pairs in refinement procedures. As Supplementary Fig. 5a demonstrates, compared with Supplementary Fig. 5b, there are many long-distance isolated base pairs marked in red, which are irrational and unstable, distorting the local structure of the loop region. After refinement, these isolated base pairs are removed.

**Fig. 6:**
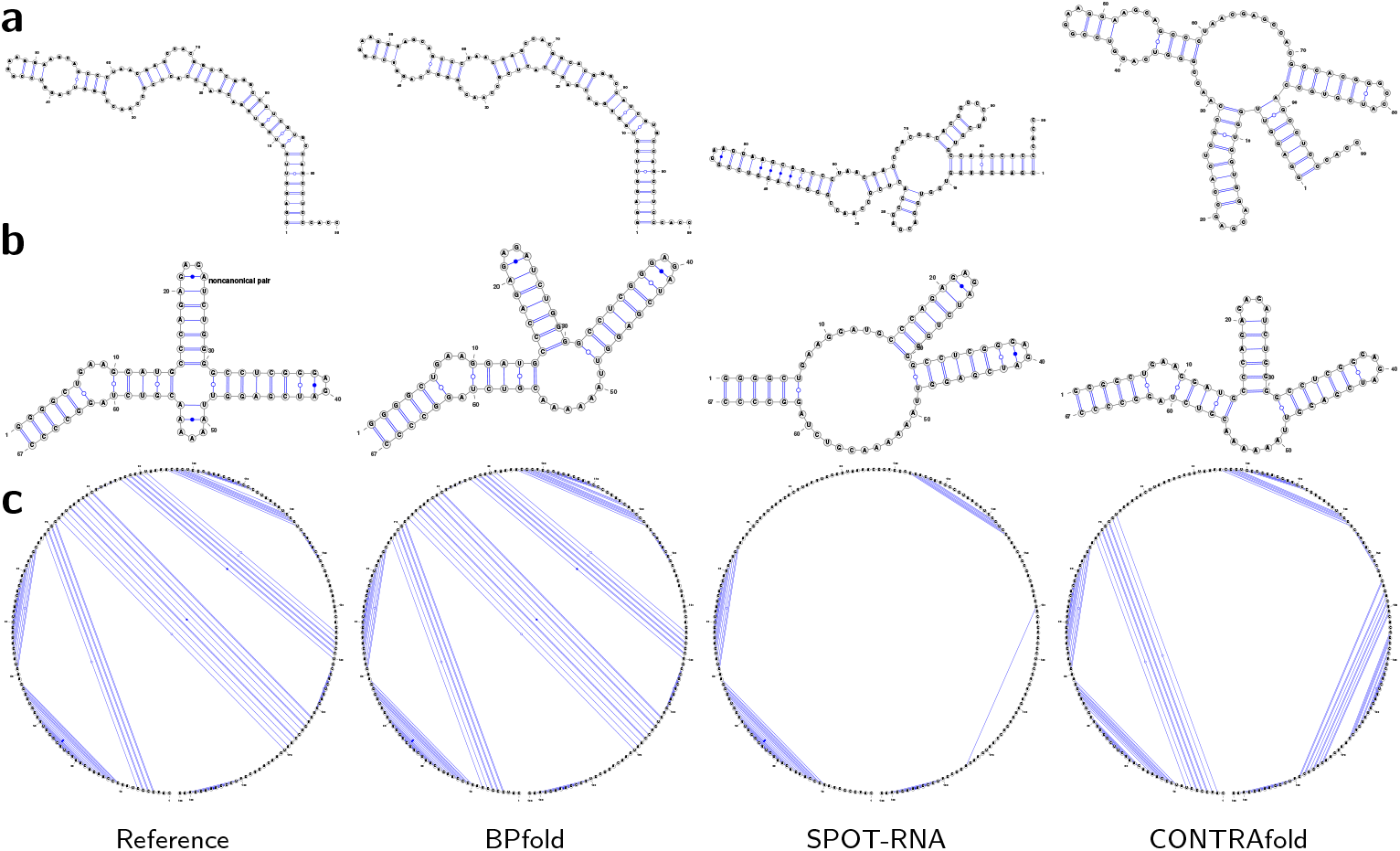
Visualization of RNA secondary structure examples predicted by BPfold along with deep learning method SPOT-RNA [36] and traditional method CONTRAfold [19]. **a** An example of brucella abortus S19 signal recognition particle from ArchiveII [49] dataset (srp Bruc.suis. AE014291), displayed in radiate style. From left to right, these structures are from native, BPfold, SPOT-RNA, and CONTRAfold, respectively. BPfold predicts the most accurate interactions compared with native reference structure, with 97% precision and 94% recall, respectively. **b** The solution structure of the delta-J-delta-K domain of EMCV IRES [60] from PDB [51] dataset (PDB ID=2NC1), displayed in radiate style, where lines with blue solid circle denote non-canonical pairs. From left to right, these structures are from native, BPfold, SPOT-RNA, and CONTRAfold, respectively. BPfold can correctly predict canonical pairs and non-canonical pairs, with 100% precision and 93% recall, respectively. **c** The crystal structures of the lariat capping ribozyme [61] from PDB [51] dataset (PDB ID=4P95), displayed in circular style. In such a dense prediction situation, BPfold predicts the most accurate base interactions than other methods, with 93% precision and 96% recall, respectively.

### 2.7 Constructing confidence index for reliable prediction

To measure the reliability of the predicted secondary structures, we further construct a confidence index, which supplies a quality assessment of predicted secondary structures in case the native structures are not available. In BPfold, the neural network directly generates the contact map 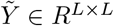, then we apply structural constraints on 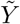 in refineme procedures, obtaining the final contact map 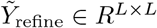. Therefore, the difference between 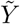 and 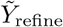 reflects the reliability and quality of predicted structures to some extent, which means that the less difference between 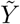 and 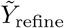 indicates the more reliability and accuracy of the output contact map that meets the structural constraints. As a result, to form the confidence index, we compute the cosine similarity between 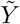 and 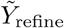 and scale it to the range of [0, 1], which can be formulated as:

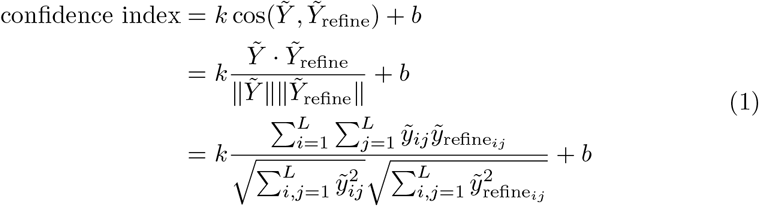

where *k* = 1.522 and *b* = −0.086 are empirically selected coefficients for adjusting the range of confidence index to [0, 1] according to predicted structures from all currently available test datasets. For robust output, the final confidence index will further be truncated to 0 or 1 if it exceeds the range of [0, 1].

To demonstrate the validity of the proposed confidence index, we compute the two-sided Pearson’s correlation coefficient between confidence indexes and F1 scores on five datasets (*i*.*e*., bpRNA-TS0, bpRNA-new, ArchiveII, Rfam12.3-14.10, PDB) together with mixed total datasets which consisting of 16,178 different RNAs. As Supplementary Table 5 shows, we achieve high correlation coefficients of 0.641, 0.676, 0.728, 0.692, 0.675, and 0.658 with low p-values, respectively, which indicates the correlation between confidence index and F1 score and further suggests that our proposed confidence index provides us a referable and reliable insight of the predicted RNA secondary structures. Fig. 7 and Supplementary Fig. 6 display direct views of the strong correlation between the designed confidence index and the F1 score metric on different datasets.

**Fig. 7:**
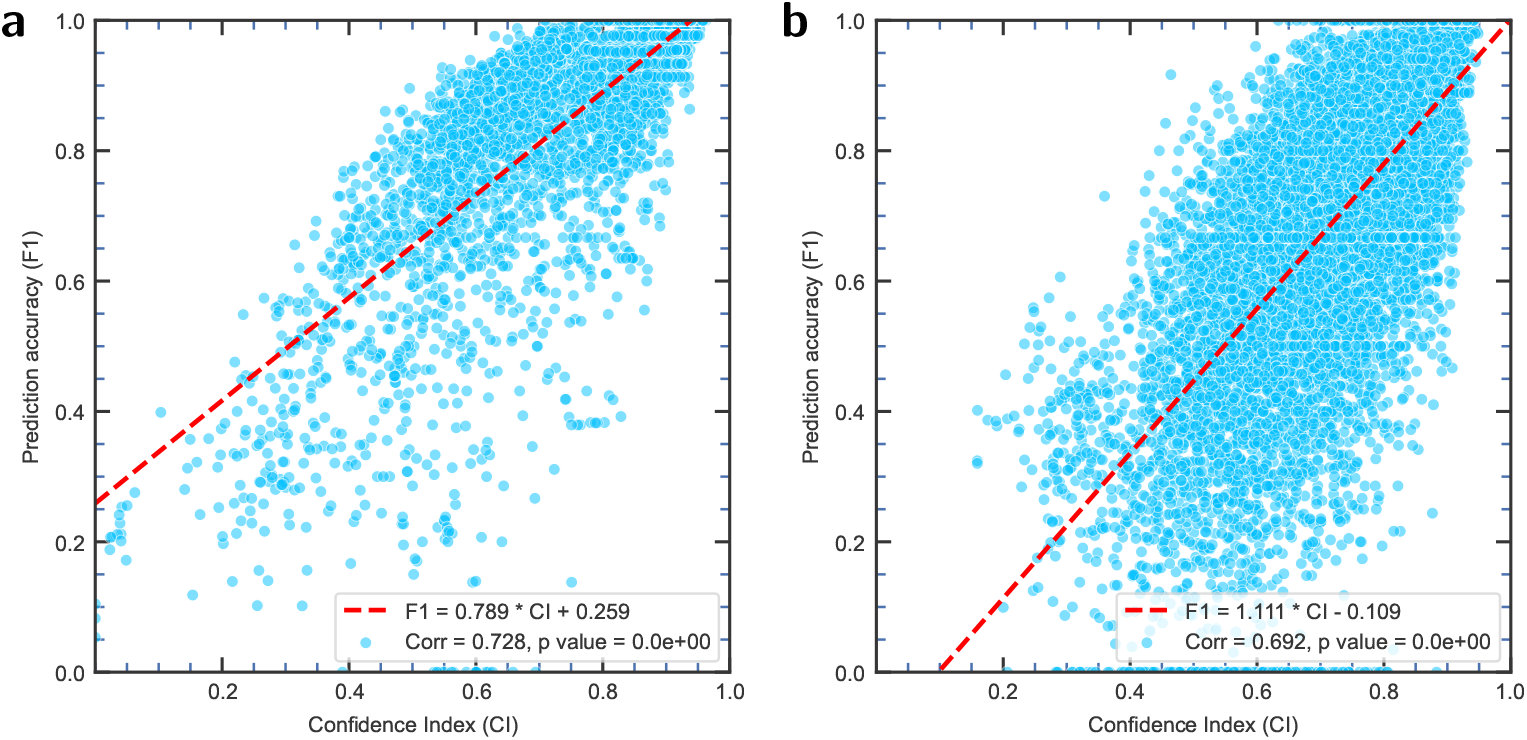
Two-sided Pearson’s correlation between F1 score and the estimated confidence index on archiveII and Rfam12.3-14.10 datasets. We compute the cosine similarity between the contact map generated from the neural network and the contact map after refinement. The Pearson correlation coefficient between the prediction accuracy (F1 score) and confidence index reaches 0.728 and 0.692, respectively. The approximate relation between F1 and CI, together with the correlation coefficients are displayed in the bottom right corner of each figure. **a** archiveII (n=3,966 RNAs), with 95% confidence interval=[0.713, 0.742]. **b** Rfam12.3-14.10 (n=10,791 RNAs), with 95% confidence interval=[0.682, 0.702].

## 3 Discussion

Our method, BPfold, aiming at improving the generalizability and accuracy of deep learning approaches, creatively proposes the complete space of a 3-neighbor base pair motif and its thermodynamic energy to tackle the problem of data insufficiency and out-of-distribution situation at the base-pair level and further bridges the knowledge prior with input RNA sequence by the elaborately designed base pair attention neural network block. The designed transformer network architecture and energy map representation facilitate the identification of long-distance interactions and the extension to unseen long RNA sequences. Experimental results on sequence-wise datasets and family-wise datasets confirm the superiority of BPfold over other methods in accuracy, generalizability, robustness, and inference speed. Additionally, we construct a confidence index for BPfold to provide a reference for the reliability of predicted RNA secondary structures. BPfold is publicly available and serves as an effective tool for RNA structure modeling.

More importantly, the proposed base pair motif and the idea of combining thermodynamic prior with input RNA sequences can be integrated into any other data-driven models. For instance, UFold [35] processes an image-like representation of RNA sequences and an alternative matrix representation [42] in consideration of possible hydrogen bonds of canonical base pairs, in which this matrix representation can be readily replaced by the energy map of base pair motif presented in this study. By doing so, the neural network achieves a complete perception of the interactions of a 3-neighbor base pair, much more accurate than the roughly counted number of hydrogen bonds. Besides, the computed energy scores of base pair motifs by the BRIQ [52] force field supplies a reliable estimation of base pair interactions, which works effectively in other thermodynamic energy-based methods. In fact, EternaFold [25] obtains thermodynamic measures via high-throughput experiments to update the parameters in CONTRAfold [19]. Similarly, it is possible to apply the framework of CONTRAfold with the statistical energy scores of base pair motifs. Note that this set of base pair motif energy scores can be further updated with the development of BRIQ or other tertiary structure *de novo* modeling methods.

Despite the innovations of BPfold in RNA sequence analysis, BPfold confronts several challenges. Firstly, on the one hand, the base pair motif can be further extended and more reliable by computing more neighboring bases, such as four or more upstream and downstream bases. In this study, we only model the tertiary structures of 3neighbor base pair motifs using BRIQ [52] due to the large computation cost. On the other hand, we could introduce more knowledge prior in extra forms rather than base pair motifs, such as RNA family information and SHAPE [11] data, to make a better understanding of the input RNA sequences and further enhance the generalization and accuracy of the models. Secondly, existing RNA secondary structure datasets consist of RNA sequences whose lengths are mainly less than 600 nucleotides or shorter, and existing deep learning methods, including BPfold, are also trained on these sequences. Therefore, to relieve the performance degradation brought by long sequences, BPfold applies a dynamic positional embedding for scalable feature embedding in the base pair attention block. However, the long sequence problem still remains unsolved; we believe that the key is to enrich the distribution of long sequences in the current training dataset. Thirdly, the prediction of non-canonical pairs is difficult for datadriven methods since there are few annotations (only in the PDB dataset) of these interactions for deep learning approaches to learn from. Previous DL methods [35–38] did not design a specific strategy or module to predict non-canonical pairs. As a result, the best F1-score of these methods is pretty low, only reaching 0.22 as benchmarked in [33]. Regardless of that, BPfold has the ability to predict non-canonical pairs and pseudo knots (Exemplified in Fig. 6), and the accuracy of BPfold for these interactions would be greatly improved when corresponding data annotations for training are adequate in the future. Further work may apply few-shot learning, domain-adaption, semi-supervised learning, and data augmentation to tackle this problem progressively.

The generalizability of deep learning models on newly discovered unseen RNA families is an inevitable issue in current research. Technically speaking, there are various common techniques to relieve the overfitting of models. As for training, we can apply early stopping to stop the learning of models before the model tends to overfitting on training data. Besides, as for the model, we also apply multiple-fold cross-validation to decrease the systematic error and select the best hyper-parameters of model architecture according to the amount of data. In this study, we innovatively propose the base pair motif and energy map from the data aspect to fully cover the data distribution at the base-pair level. BPfold receives the information not only from the RNA sequence but also the energy map of all paired canonical pairs in this RNA sequence, which allows the specific prediction of each input RNA sequence and hinders the overfitting of similar RNA sequences. As a result, evidenced in experiments on family-wise dataset Rfam12.3-14.10, BPfold achieves the best performances against other methods, revealing the great generalizability and accuracy in modeling RNA secondary structures from unseen RNA families. Apart from base pair motif, highthroughput chemical probing data [25, 63] such as SHAPE [64] can be applied to provide information of the chemical activity of nucleotide bases. Note that because the auxiliary information involved in existing deep learning methods such as BPfold, MXfold2, UFold, and SPOT-RNA is implicitly learned by neural networks, further improvements in modeling RNA secondary structures can be achieved when learning approaches can explicitly integrate physical laws into neural networks.

In summary, we show a great prospect of BPfold in improving generalizability and accuracy with base pair motif for RNA secondary structure prediction, expecting that BPfold will inspire further advancements in the integration of physical priors with deep learning techniques, as well as enhance our understanding of RNA structures and their biological functions.

## 4 Methods

### 4.1 Base pair motif energy as thermodynamic prior

The performance and generalizability of deep learning models are highly dependent on training data, which currently may be hampered by the lack of structural data and the limitation of data diversity. To tackle this, we aim at improving the coverage of data at the base-pair level and bringing thermodynamic energy to RNA sequence modeling. In view of the locality of the secondary structures of RNA, we define the *base pair motif*, a canonical base pair (i.e., A-U, U-A, G-C, C-G, G-U, and U-G) together with the *r* upstream and downstream neighboring bases of each base.

Specifically, base pair motifs can be divided into three categories, for any two bases indexed as *i* and *j*(*i < j*) of an RNA sequence: (1) Inner hairpin base pair motif (BPM_iH_): While neighboring bases extend to inner sequence and *j* − *i* ≤2*r*, the downstream of the base *i* and the upstream of base *j* are continuous in sequence and form a hairpin loop, with the base pair motif denoted as [*i, i* + 1, …, *j*]. There are 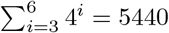 sequences for each canonical base pair of this category. (2) Inner chainbreak base pair motif (BPM_iCB_): While neighboring bases extend to inner sequence and *j* − *i >* 2*r*, the downstream of base *i* is not continuous with the upstream of base *j*, forming a chain-break. Therefore, the base pair motif consists of two chains, denoted as [*i, i* + 1, …, *i* + *r* : *j* − *r, j* − *r* + 1, …, *j*], where : represents chain-break of RNA sequence. There are 4^6^ = 4096 sequences for each canonical base pair of this category. (3)Outer base pair motif (BPM_oCB_): While neighboring bases extend to outer both ends of the sequence, the base pair motif is denoted as [*j, j* + 1, …, *j* + *r* : *I* − *r, I* − *r* + 1, …, *i*]. Besides that, we also deal with special corner cases while base *i* or base *j* has no sufficient *r* neighboring bases upstream or downstream. There are 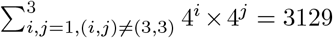 sequences for each canonical base pair of this category.In total, there are 6 × (5440 + 4096 + 3129) = 75990 base pair motifs for all six kinds of canonical base pairs (i.e., A-U, U-A, G-C, C-G, G-U, and U-G). In the formation of these, the paired bases are always located at the beginning and the end of each category of base pair motif.

After enumerating all possible sequences of *r*-neighbor base pair motifs, we use our previous *de novo* RNA tertiary structures modeling method BRIQ [52] to model the tertiary structure of each base pair motif and extract energy score *E*_bpm_(*L*_bpm_, *I*_break_) estimated by BRIQ force field of whole motif sequence, where *L*_bpm_ is the length of base pair motif and *I*_break_ represents whether there is a chain-break in this motif. Then we compute the energy score *E*_bpm_ of each motif by eliminating the influence of a single strand and normalizing the value with the maximum energy of the motif with the same length in the same category, which can be formulated as:

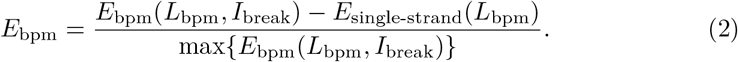

Finally, we establish a base pair motif library with these energy items and tertiary structures. As Fig. 2 demonstrates, with this energy table, for any two bases of RNA sequence with index being *i* and *j*(*i < j*), we query the outer/inner base pair motif energy items of corresponding outer/inner base pair motif for any canonical base pair (i.e., A-U, U-A, G-C, C-G, G-U and U-G) and we set the energy of any other noncanonical pair to zero. Therefore, for an RNA sequence with *L* nucleotides, we can obtain the outer/inner energy maps *M*^*µ*^ and *M*^*ν*^ in the shape of *L* × *L*.

### 4.2 Deep neural network with base pair attention

As shown in Figure 1, BPfold is a deep neural network consisting of consecutive *N* modified transformer blocks. In each transformer block, there is an elaborately designed base pair attention block (Fig. 1b), composed of hybrid convolutional block (Fig. 1c) which applies the squeeze-and-excitation (SE) block [65](Fig. 1d) to adaptively re-calibrate the channel-wise feature response and explicitly build the relationship of energy maps, along with an enhanced self-attention mechanism that aggregates attention map from RNA sequence and base pair motif energy, learning thermodynamic knowledge in the complete space of *r*-neighbor base pair motif to improve the generalizability of model in situation of unseen RNA sequence and families.

The original transformer block [47] is composed of a multi-head self-attention module (MSA), followed by a feed-forward network (FFN) which consists of a two-layer multi-layer perceptron (MLP) with a GELU activation. A LayerNorm (LN) layer is adopted before each MSA and each FFN, and a residual shortcut is adopted after each module.

To integrate the thermodynamic priors in the form of energy maps into this attention mechanism, we design the base pair attention block. As illustrated in Figure 1b, when processing self-attention, a base pair attention block applies several 3 × 3 convolutional layers (denoted as *CONV*) to the energy map to establish the relationship among base pairs. Furthermore, This attention block adds the thermodynamic feature map to the attention map of sequence features, imposing the thermodynamic relationship of base pairs on RNA sequences. More specifically, the input of the neural network is an RNA sequence of length *L*, containing four bases, *i*.*e*., A, C, G, U (other unknown bases will be converted to the above four bases). The input RNA sequences are firstly padded with “START”, “END” and “EMPTY” tokens to a uniform length *L*_max_ to deal with variable lengths, which are then encoded into a *D*-dimensional embedding using trainable parameters, forming the input feature *X* in a shape of *L*_max_ × *D*. Meanwhile, the energy maps *M*^*µ*^ and *M*^*ν*^ of the outer base pair motif and inner base pair motif are prepared according to the input RNA sequence and the stored energy table. Similar to the input feature of RNA sequences, both *M*^*µ*^ and *M*^*ν*^ energy maps are padded with zero to the shape of *L*_max_ × *L*_max_. The base pair attention block processes can be formulated as follows: (1) Obtaining base pair attention maps:

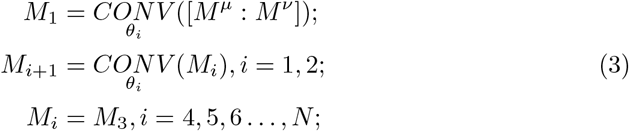

where *M*_*i*_ is the *i*-th base pair attention block, [*M*^*µ*^ : *M*^*ν*^] denotes the concatenation of base pair energy maps *M*^*µ*^ and *M*^*ν*^, and *θ* represents learnable parameters. (2) Integrating base pair attention maps with sequence feature maps into the transformer block:

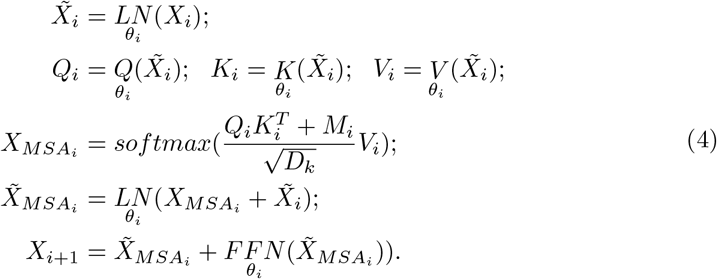

where *i* = 1, 2, …, *N* with *N* being the number of transformer blocks. After *N* base pair attention blocks, we obtain the output orthogonal matrix 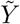 in shape of *L*_max_ × *L*_max_ by applying matrix multiplication between *X*_*N*_ and its transpose 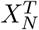, namely 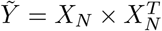, which represents the possibility score of each nucleotide being paired with other nucleotides.

### 4.3 Training strategy and structure refinement

To relieve the impact of efficiency brought by padding, we apply a length-matching strategy for sampling a mini-batch at the training stage. Specifically, we set a series of buckets {*B*_0_, *B*_1_, *B*_2_, …} and assign each input RNA sequence to a bucket 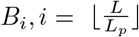 where *L* is the length of RNA sequence and *L*_*p*_ is the predefined interval of buckets. Mini-batches are sampled from the same bucket, which leads to a controllable padding size *L*_*p*_. This length-matching strategy is especially effective in dealing with the varying lengths of input RNA sequences from tens for short sequences to thousands for large sequences.

BPfold is implemented in PyTorch framework [66] and trained by minimizing the binary cross-entropy between the predicted contact matrix 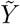 and the true contact matrix *Y* using the ADAM optimization algorithm. The number of parameters is listed in Supplementary Table 6. To leverage the imbalanced distribution of paired bases and unpaired bases, we adopt a positive weight *ω* = 300 to derive the loss function as below:

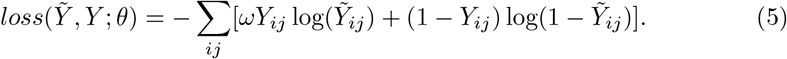

where *θ* denotes all learnable parameters of the neural network, *I* ∈ {1, 2, …, *L*} and *J* ∈ {1, 2, …, *L*} denote the row and column, respectively, index of matrices.

When predicting an RNA sequence, we apply refinement procedures to the output contact map 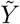 generated by the BPfold neural network to impose physical constraints on RNA secondary structures and rule out invalid base pairs. Specifically, the following rules of constraints are considered in which the first three rules are inspired from previous study [35, 38]: (1) Only Waston-Crick pairs (*i*.*e*., A-U, U-A, G-C, C-G) and Wobble pair (G-U, U-G) are allowed for canonical pairs while others are allowed for non-canonical base pairs; (2) A loop region has at least two bases for ruling out sharp loops; (3) Overlapping pairs are discarded. We encode these constraints as matrix transformations and apply them to the output contact matrix. Since the output contact matrix is already a symmetric matrix 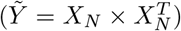, we do not explicitly declare symmetric processing. (4) Isolated base pairs are removed. An isolated base pair has no consecutive neighboring helix and is not stable enough to form a base pair in most situations. We verify all paired bases and remove isolated base pairs to rule out long-distance unstable interactions.

### 4.4 Datasets and evaluation

We utilize several widely used open-source benchmark datasets for evaluating the performances of our proposed BPfold and comparing it with state-of-the-art RNA secondary structure prediction methods. Specifically, these benchmark datasets are as follows:

- RNAStralign [55]: This dataset contains 37,149 RNA sequences from eight RNA families, with sequence lengths ranging from approximately 30 to 3,000 nucleotides (nt). Similar to previous work [35, 38, 39], we remove redundant sequences and invalid secondary structures, and obtain a total of 29,647 unique sequences. Furthermore, we filter sequences with lengths no more than 600 nt, forming a training dataset that consists of 19,313 sequences.
- bpRNA-1m [50]: This dataset contains 102,318 RNA sequences from 2,588 RNA families. Following MXfold2 [39], we use CD-HIT program [59] to remove similar sequences with a cut-off of 80% and split the processed dataset into two sub-dataset for training and testing, named TR0 and TS0, which contain 12,114 and 1,305 sequences respectively, with sequence lengths ranging from 22 to 499 nt.
- ArchiveII [49]: This dataset is the most widely used benchmark dataset for evaluation of RNA secondary structures, consisting of 3,966 RNA sequences from ten RNA families, *i*.*e*., 5s rRNA 16s rRNA, 23s rRNA, tRNA, tmRNA, telomerase RNA RNase P, SRP, group I Intron, and group II Intron. Among them, 3,911 sequences have a length below 600 nt while the other 55 RNA sequences are all from group II Intron which have a max length of 1,800 nt.
- Rfam12.3-14.10: We construct this dataset by initially collecting 50,779 RNA sequences that are newly added to the latest version of Rfam [17, 18], namely from Rfam version 12.3 to Rfam version 14.10, which includes newly added cross-family sequences that are not present in the bpRNA training dataset. After using CD-HIT-EST [59] to remove redundant sequences with sequence similarity of more than 80%, this dataset contains 10,791 unique sequences from 1,992 RNA families, with lengths of 68 sequences ranging from 600 nt to 951 nt and the other ranging from 26 nt to 600 nt. We employ this dataset for family-wise evaluation. We also evaluate models on bpRNA-new dataset derived from Rfam version 12.3 to Rfam version 14.2 by MXfold2 [39] which contains 5,401 sequences, with sequence length ranging from 33 nt to 489 nt.
- PDB [51]: This dataset is a benchmark dataset, consisting of 116 RNA sequences with high-resolution (*<* 3.5Å) RNA X-ray structures, with sequence lengths ranging from 32 to 355 nt. According to previous study [35–37], the PDB dataset is divided into three sub datasets, i.e., TS1, TS2, and TS3, with 60, 38, and 18 sequences, respectively.

For performance evaluation of predicted RNA secondary structures, we use precision (P), recall (R, a.k.a. sensitivity), F1 score, and interaction network fidelity (INF) to assess the quality of base pair prediction, which is a binary classification problem. We calculate the macro-averages of these metrics of canonical base pairs. Specifically, these metrics are defined as below:

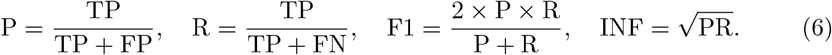

where TP, FP, and FN denote true positive (the number of correctly predicted base pairs), false positive (the number of incorrectly predicted base pairs), and false negative (the number of base pairs whose reference structures are not predicted), respectively.

## Data availability

All data used in this study are available at Zenodo [67] and GitHub (https://github.com/heqin-zhu/BPfold/releases). These data include datasets, base pair motif energy scores, model parameters and source data. The source data underlying Tables 1, 2, S2-S5, and Figs. 4, 5, 7, S3, S4, S6 are provided in the Source Data file. Source data are provided with this paper.

## Code availability

The source code and program tool of BPfold method is publicly available at Zenodo [67] and GitHub (https://github.com/heqin-zhu/BPfold/releases).

## Acknowledgements

This work is supported by National Natural Science Foundation of China (62271465 to S.K.Z., 32370581 to P.X.), Open Fund Project of Guangdong Academy of Medical Sciences, China (YKY-KF202206 to S.K.Z.), and Suzhou Basic Research Program (SYG202338 to S.K.Z.).

## Author contribution

H.Z. constructed the motif library, designed the network architectures, conducted the experiments, analyzed the results, and wrote the paper. P.X. and S.K.Z. conceived the idea, supervised the study, and designed the experiments. P.X. also participated in the design of the core components. F.T. carried out part of the experiments and participated in the design of networks. Q.Q. and K.C. helped in the analysis of data results. All authors read, contributed to the discussion, and approved the final paper.

## Additional information

## Supplementary information

The supplemesntary for this paper is publicly available.

