## Supplementary for "Deep generalizable prediction of RNA secondary structure via base pair motif energy"

### Supplementary Materials for Deep generalizable prediction of RNA secondary structure via base pair motif energy

### 1 Supplementary figures

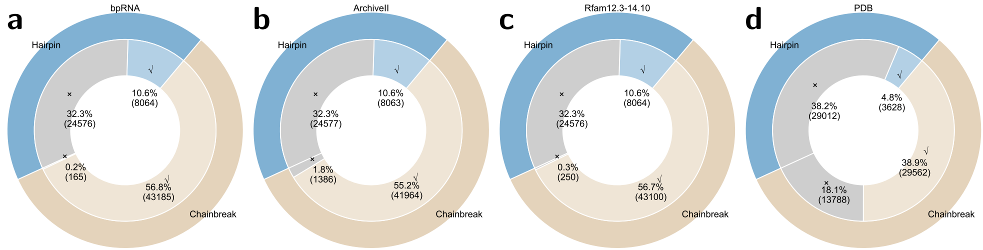


**Supplementary Fig. 1: Visualization of the data coverage of hairpin and chainbreak base pair motifs in current datasets.** The outer ring of each pie chart represents the hairpin and chainbreak distribution in the whole base pair motif library (75,990 motifs, blue for hairpin motifs, orange for chainbreak motifs). The inner ring represents the hairpin and chainbreak distribution in each dataset (blue for hairpin motifs, orange for chainbreak motifs, and grey for missing motifs). **a** bpRNA (n=1,305 RNAs) [1]. **b** ArchiveII (n=3,966 RNAs) [2]. **c** Rfam12.3-14.10 (n=10,791 RNAs) [3, 4]. **d** PDB (n=116 RNAs) [5].

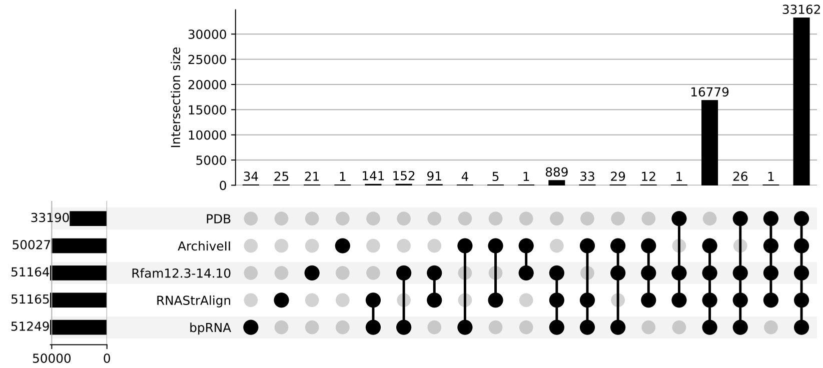


**Supplementary Fig. 2: Upset illustrations of the interaction of base pair motifs from various datasets**.

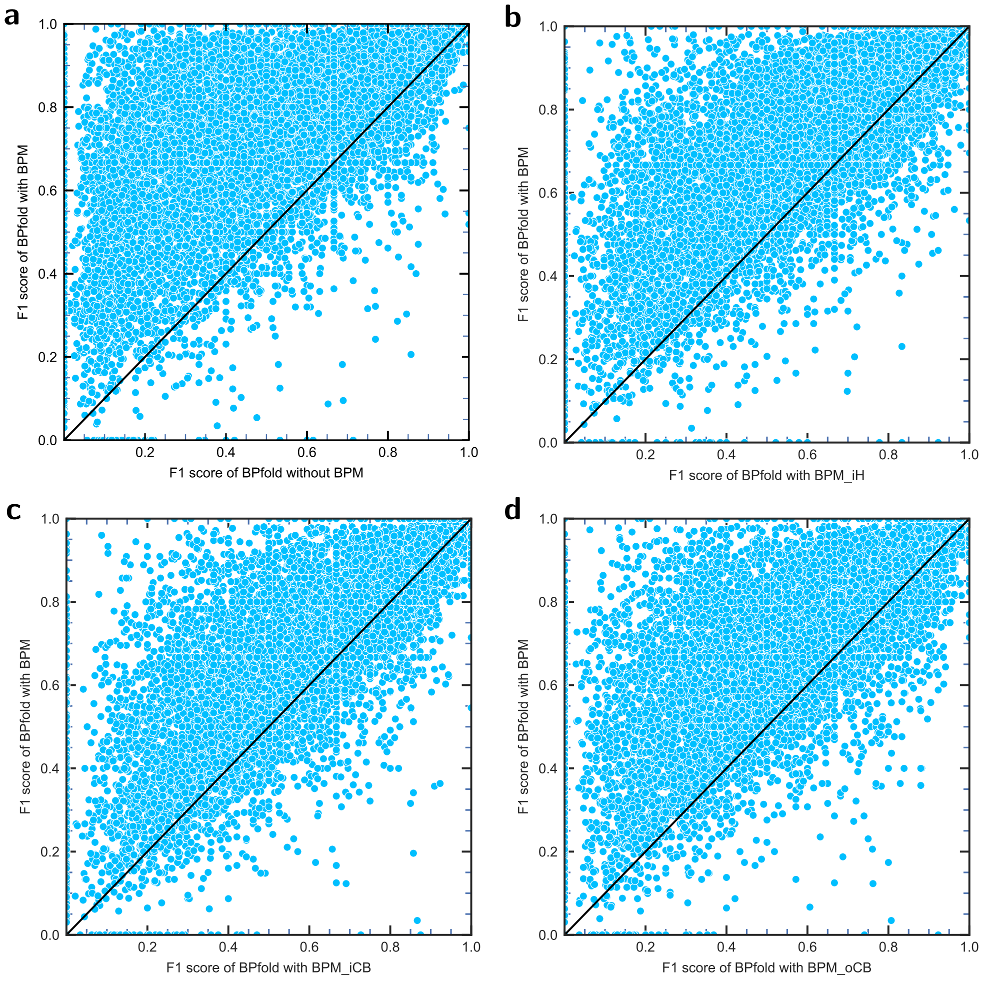


**Supplementary Fig. 3**: **Head-to-head comparison of BPfold with BPM against other configuration on Rfam12.3-14.10 (n=10,791 RNAs)** [3, 4] **a** without BPM. **b** with $BPM_{iH}$. **c** with $BPM_{iCB}$. **d** with $BPM_{oCB}$.

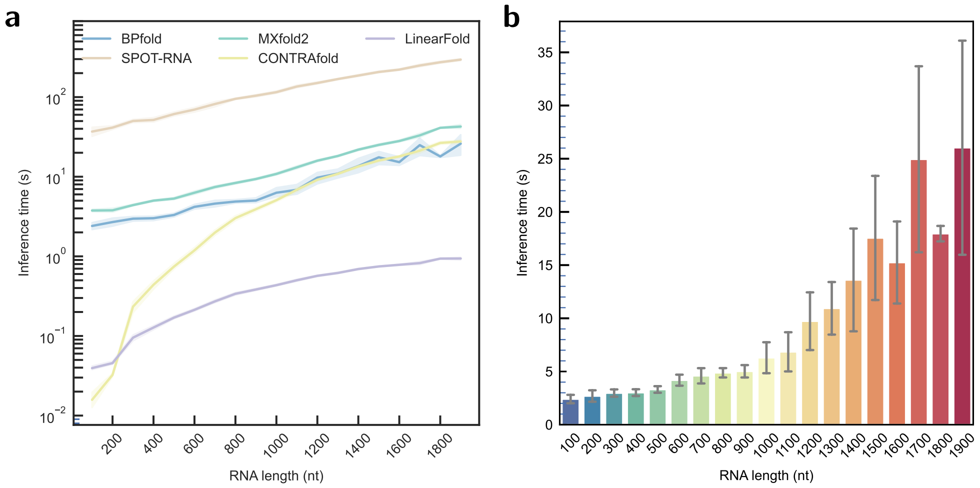


**Supplementary Fig. 4**: **Evaluation of the inference time of BPfold and other methods on a selected dataset which contains 174 RNAs with lengths varying from 60 to 1,851 nucleotides uniformly.** To decrease the influence of randomness, we repeat 10 times to predict each RNA sequence. **a** Comparison with other methods. **b** Data are presented as mean values ± SD. BPfold predicts RNA secondary structures robustly and fast, which takes less than 10 seconds for RNAs no longer than 1,000 nucleotides, and less than 40 seconds for RNAs no longer than 1,851 nucleotides.

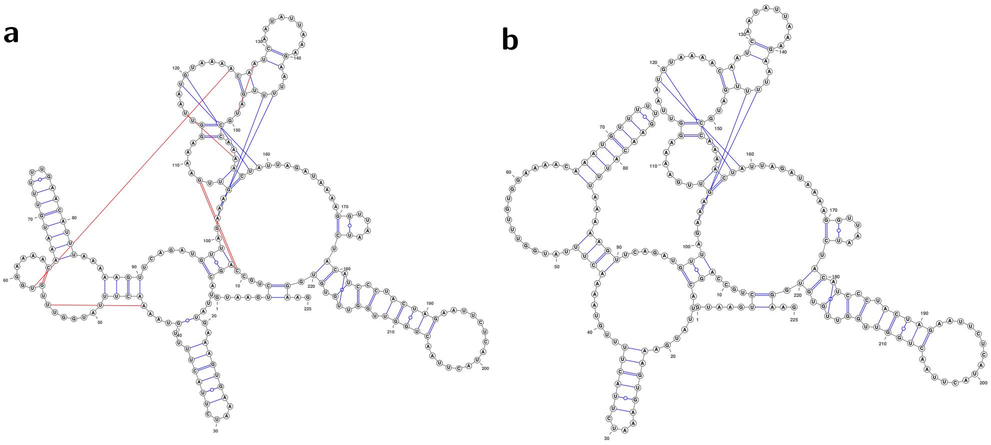


**Supplementary Fig. 5**: **Visualization of example RNA secondary structures predicted by BPfold with and without removing isolated base pairs. a** The secondary structure of an RNA sequence predicted by BPfold without removing isolated base pairs. **b** Compared with subfigure **a**, this secondary structure is further processed from the structure in **a** by removing isolated base pairs to eliminate the influence of long-distance unstable interactions.

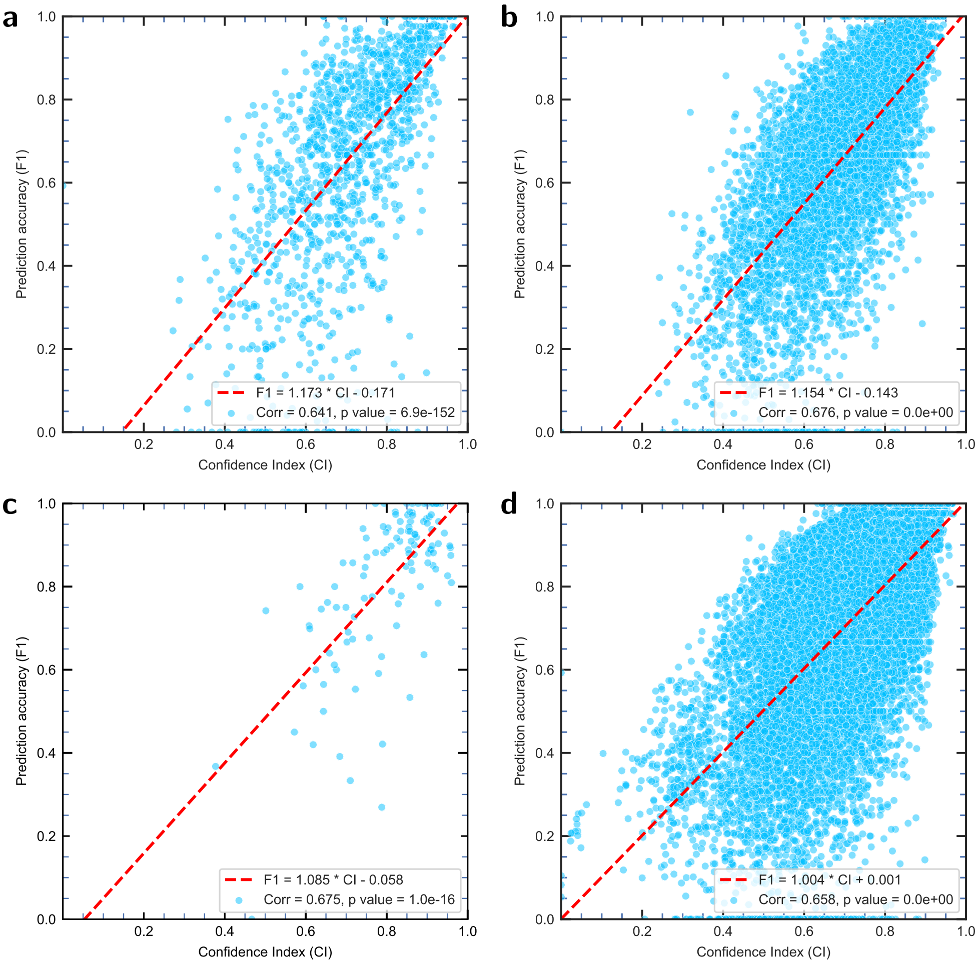


**Supplementary Fig. 6**: **Two-sided Pearson's correlation between F1 score and the estimated confidence index on bpRNA, bpRNA-new, PDB, and total datasets.** We compute the cosine similarity between the contact map generated from the neural network and the contact map after refinement. The Pearson correlation coefficient between the prediction accuracy (F1 score) and confidence index reaches 0.641, 0.676, 0.675 and 0.658, respectively. The approximate relation between F1 and CI, together with the correlation coefficients are displayed in the bottom right corner of each figure. **a** bpRNA (n=1,305 RNAs), with 95% confidence interval=[0.608, 0.672]. **b** bpRNA-new (n=5,401 RNAs), with 95% confidence interval=[0.662, 0.690]. **c** PDB (n=116 RNAs), with 95% confidence interval=[0.562, 0.763]. **d** The total datasets contains different RNAs from Rfam, ArchiveII, and PDB (n=16,178 RNAs), with 95% confidence interval=[0.649, 0.667].

#### 2 Supplementary tables

**Supplementary Table 1**: Summary of training and test datasets in our experiments.

| Dataset | phase | # sequences | length |
|---|---|---|---|
| RNAStrAlign [6] | train | 19,313 | 30-600 |
| bpRNA-1m [1]: TR0, VL0 | train | 12,114 | 33-498 |
| bpRNA-1m [1]: TS0 | test | 1,305 | 22-499 |
| ArchiveII [2] | test | 3,966 | 28-1,800 |
| Rfam12.3-14.10 [3, 4] | test | 10,791 | 26-951 |
| bpRNA-new (Rfam12.3-14.2) | test | 5,401 | 33-489 |
| PDB [5] | train | 652 | 6-422 |
| PDB [5] | test | 116 | 32-355 |

**Supplementary Table 2**: Ablation study of BPfold under five different configurations of base pair motif energy on familly-wise dataset Rfam12.3-14.10.

| Configuration | Rfam12.3-14.10 | | | |
|---|---|---|---|---|
| | INF | F1 | Precision | Recall |
| w/ BPM | 0.694 | 0.689 | 0.660 | 0.741 |
| w/ $BPM_{iH}$ | 0.561 | 0.555 | 0.563 | 0.569 |
| w/ $BPM_{iCB}$ | 0.598 | 0.592 | 0.577 | 0.630 |
| w/ $BPM_{oCB}$ | 0.556 | 0.552 | 0.549 | 0.574 |
| w/o BPM | 0.445 | 0.438 | 0.480 | 0.426 |

**Supplementary Table 3**: Performance evaluation of RNA secondary structure prediction approaches on bpRNA-new (5,401 RNAs, from Rfam12.3-14.2) and PDB subsets TS1 (60 RNAs).

| Method | bpRNA-new | | | | PDB-TS1 | | | |
|---|---|---|---|---|---|---|---|---|
| | INF | F1 | Precision | Recall | INF | F1 | Precision | Recall |
| BPfold | 0.655 | 0.647 | 0.604 | 0.723 | 0.760 | 0.756 | 0.788 | 0.740 |
| SPOT-RNA | 0.623 | 0.617 | 0.620 | 0.641 | 0.785 | 0.778 | 0.847 | 0.738 |
| MXfold2 | 0.640 | 0.632 | 0.585 | 0.710 | 0.728 | 0.723 | 0.782 | 0.683 |
| ContextFold | 0.585 | 0.579 | 0.546 | 0.635 | 0.715 | 0.709 | 0.760 | 0.682 |
| ContraFold | 0.634 | 0.627 | 0.608 | 0.680 | 0.697 | 0.692 | 0.770 | 0.653 |
| EternaFold | 0.643 | 0.633 | 0.569 | 0.736 | 0.729 | 0.726 | 0.752 | 0.712 |
| LinearFold | 0.625 | 0.617 | 0.649 | 0.645 | 0.669 | 0.663 | 0.764 | 0.621 |
| RNAfold | 0.626 | 0.617 | 0.552 | 0.720 | 0.679 | 0.676 | 0.697 | 0.666 |
| SimFold | 0.605 | 0.596 | 0.536 | 0.692 | 0.639 | 0.636 | 0.664 | 0.622 |
| RNAStructure | 0.614 | 0.604 | 0.539 | 0.711 | 0.677 | 0.674 | 0.695 | 0.665 |

**Supplementary Table 4**: Performance evaluation of RNA secondary structure prediction approaches on PDB subsets, TS2 (38 RNAs) and TS3 (18 RNAs).

| Method | PDB-TS2 | | | | PDB-TS3 | | | |
|---|---|---|---|---|---|---|---|---|
| | INF | F1 | Precision | Recall | INF | F1 | Precision | Recall |
| BPfold | 0.934 | 0.933 | 0.938 | 0.934 | 0.760 | 0.757 | 0.803 | 0.724 |
| SPOT-RNA | 0.884 | 0.881 | 0.911 | 0.861 | 0.762 | 0.754 | 0.841 | 0.698 |
| MXfold2 | 0.869 | 0.864 | 0.933 | 0.817 | 0.781 | 0.775 | 0.851 | 0.724 |
| ContextFold | 0.828 | 0.823 | 0.894 | 0.773 | 0.654 | 0.651 | 0.701 | 0.614 |
| ContraFold | 0.848 | 0.844 | 0.892 | 0.811 | 0.742 | 0.735 | 0.826 | 0.674 |
| EternaFold | 0.843 | 0.840 | 0.863 | 0.828 | 0.692 | 0.690 | 0.734 | 0.655 |
| LinearFold | 0.828 | 0.821 | 0.889 | 0.781 | 0.698 | 0.686 | 0.812 | 0.611 |
| RNAfold | 0.881 | 0.878 | 0.921 | 0.849 | 0.706 | 0.704 | 0.736 | 0.679 |
| SimFold | 0.887 | 0.884 | 0.932 | 0.848 | 0.757 | 0.755 | 0.784 | 0.734 |
| RNAStructure | 0.899 | 0.898 | 0.924 | 0.877 | 0.701 | 0.700 | 0.727 | 0.677 |

**Supplementary Table 5**: Pearson correlation between prediction accuracy (F1 score) and confidence index (CI) of BPfold on five datasets. Two-sided hypothesis, 95% confidence intervals, approximate relation of CI and F1 are displayed. The total dataset contains different RNAs from Rfam, ArchiveII, PDB.

| Dataset | Correlation | Confidence | p-value | Relation |
|---|---|---|---|---|
| ArchiveII [2] | 0.728 | $[0.713, 0.742]$ | 0.000 | $\mathrm{F1} = 0.789 * \mathrm{F1} + 0.259$ |
| bpRNA-1m [1]: TS0 | 0.641 | $[0.608, 0.672]$ | $6.937 \times 10^{-152}$ | $\mathrm{F1} = 1.173 * \mathrm{CI} - 0.170$ |
| bpRNA-new | 0.676 | $[0.662, 0.690]$ | 0.000 | $\mathrm{F1} = 1.154 * \mathrm{CI} - 0.143$ |
| Rfam12.3-14.10 [3, 4] | 0.692 | $[0.682, 0.702]$ | 0.000 | $\mathrm{F1} = 1.111 * \mathrm{CI} - 0.109$ |
| PDB [5] | 0.675 | $[0.562, 0.763]$ | $1.002 \times 10^{-16}$ | $\mathrm{F1} = 1.085 * \mathrm{CI} - 0.058$ |
| total | 0.658 | $[0.649, 0.667]$ | 0.000 | $\mathrm{F1} = 1.004 * \mathrm{CI} + 0.001$ |

**Supplementary Table 6**: Number of parameters of recent deep learning models.

| Method | # Parameter |
|---|---|
| BPfold | 7,962,416 |
| SPOT-RNA | 7,759,445 |
| MXfold2 | 47,346 |
| UFold | 8,641,377 |
| e2efold | 718,863 |
